# *In situ* Architecture of the Tad Pilus Machine in *Caulobacter crescentus*

**DOI:** 10.1101/2025.11.05.686773

**Authors:** James Iarocci, Gregory B. Whitfield, Ryu F. Williston, Michael R. Wozny, John F. Presley, Courtney K. Ellison, Yves V. Brun, Shuaiqi Guo

## Abstract

The Tight adherence (Tad) pilus is a broadly distributed and evolutionarily distinct subclass of type IV pili that mediates cell adhesion, biofilm formation, predation, and surface sensing in many bacteria, including *Caulobacter crescentus*, *Myxococcus xanthus*, *Vibrio vulnificus*, and *Bifidobacterium breve*. Tad pili undergo cycles of extension and retraction powered by a cell-envelope-embedded nanomachine. Despite their biological importance, the architecture and assembly mechanism of the Tad pilus system have remained poorly understood. Although cryo-electron tomography (cryo-ET) has elucidated the *in situ* structures of other type IV pilus systems, no intact Tad machine structure has previously been reported. Here, we use cryo-ET and subtomogram averaging to resolve the near-native architecture of the *C. crescentus* Tad pilus within the bacterial cell envelope. 3D classification further reveals multiple assembly intermediates, and integrative modelling incorporating AlphaFold3 predictions help define the spatial arrangement of all core components. The resulting structural framework gives insight into the stepwise assembly process of the *C. crescentus* Tad pilus machine. Altogether, our results provide an *in situ* architectural model of the Tad pilus machine, establishing a foundation for understanding homologous systems across a broad range of bacteria.

**Importance:** Investigating the Tad pilus nanomachine in a genetically tractable, non-pathogenic organism like *Caulobacter crescentus* provides a powerful model for elucidating the architecture and functional dynamics of this widespread system. Insights gained from studying the Tad machinery can improve our understanding of related Tad pilus systems in pathogenic bacteria such as *Aggregatibacter actinomycetemcomitans*, where Tad pili are a key determinant of biofilm formation and chronic infection. Additionally, the remarkable functional diversity of Tad systems, ranging from surface sensing in *C. crescentus* to bacterial predation in *M. xanthus*, highlights their broad biological relevance. By revealing the *in situ* structure and assembly mechanism of the Tad pilus biosynthetic machinery, this study advances our understanding of a major class of bacterial nanomachines and may thus provide structural insights that could inform the development of new therapeutic strategies targeting pilus-mediated virulence.

## Introduction

Many bacteria use surface-exposed hair-like pili (or fimbriae) to mediate critical interactions with the environment, including attachment, motility, DNA uptake, and biofilm formation^1–5^. Among these systems, type IV pili (T4P) are distinguished by their ability to undergo rapid cycles of extension and retraction, a dynamic behavior that generates substantial mechanical force^6–10^. These dynamics are driven by the multi-protein T4P nanomachine embedded in the bacterial cell envelope, which encompasses several subsystems across bacteria – the Type IVa pilus (T4aP), Type IVb pilus (T4bP), and the Type IVc pilus, also known as the tight <u>ad</u>herence (Tad) pilus^11^. These subsystems share homologous core components and exhibit similar functions, whilst differing in the architecture and composition of their biosynthetic machinery.

Of the T4P systems, the Tad pilus remains the least understood^11,12^. Initially discovered in *Aggregatibacter actinomycetemcomitans*^13,14^, phylogenetic analyses suggest that the Tad pilus originates from the archaeal branch of the type IV filament (TFF) superfamily, a broad group of filament-forming nanomachines that includes pili and secretion systems found across bacteria and archaea^15,16^. From this archaeal origin, the Tad pilus has since found its place in bacteria through horizontal gene transfer^16^. In contrast, other T4P systems originate from the bacterial branch of the TFF superfamily, and are thus less related to the Tad pilus machine, suggesting that insights from other T4P systems may not directly apply to Tad systems. For example, one key distinction that separates Tad pili from other T4P is that they always rely on a single bifunctional ATPase motor for both extension and retraction, as opposed to two monofunctional ATPases in some other systems^17,18^.

Among bacteria, Tad pili are widely distributed and regulate a variety of biological functions. They induce cell aggregation and biofilm formation in *A. actinomycetemcomitans*, promote twitching motility in *Liberibacter crescens*, enable DNA uptake and transformation in *Micrococcus luteus*, and aid commensal host colonization by *Bifidobacterium breve*^11^. They also enhance virulence and predatory behaviors in multiple pathogenic and non-pathogenic species, including *Aeromonas hydrophila*, *Pseudomonas aeruginosa*, *Vibrio vulnificus*, and *Myxococcus xanthus*^11,19^. In the well-studied model organism *Caulobacter crescentus*, Tad pili play a key role in surface sensing and in orienting cells prior to irreversible surface attachment through the initiation of holdfast synthesis^20^. Consequently, *C. crescentus* has emerged as an important, genetically tractable model system for the study of Tad pilus biology, since its pili are well characterized in the context of the cell’s developmental cycle and polarity, while its Tad machine components are highly conserved across the TFF superfamily^21,22^. The core components of the *C. crescentus* Tad pilus machine, hereafter referred to as *Cc*Tad, are encoded by the *cpa* (*Caulobacter* pilus-associated) gene cluster, homologous and functionally equivalent to the *tad/rcp* gene clusters described in other bacteria using an alternative nomenclature^11,23^.

Previous studies have identified the proposed subunits of the multi-layered, cell envelope-spanning *Cc*Tad machine based on genetic and biochemical analyses^23,24^. In the outer membrane (OM), the secretin CpaC is predicted to form a pore complex through which the pilus is extended and retracted, while the functions of the putative pilotin CpaO and the lipoprotein CpaD remain unknown. In the Tad machine of *P. aeruginosa*, the homologous secretin RcpA has been shown to form a cage-like structure with 12-14-fold symmetry in the OM, aided by the pilotin TadD, which regulates its assembly and localization^11,25^. The *Cc*Tad secretin CpaC shares 40% sequence identity with *P. aeruginosa* RcpA, suggesting that a similar architecture may be present in the *Cc*Tad system. Meanwhile, in the periplasm, the *Cc*Tad system comprises CpaB and the smaller protein CpaI, which may be important to promote pilus alignment, a function observed in *P. aeruginosa* RcpC, a CpaB homolog^12^. Finally, the activity of the *Cc*Tad complex is driven by the cytoplasmic ATPase CpaF, which powers both assembly and disassembly of the *Cc*Tad pilus^11,17,26^. CpaF is likely recruited to the IM by the platform proteins CpaG and CpaH, and may be further regulated by CpaE, a ParA/MinD family ATPase that is predicted to associate with the IM through a conserved amphipathic helix^11^. While Cryo-EM structures of CpaF and some orthologues of CpaC and CpaB have been determined^12,17,25,26^, the architecture of the full Tad pilus machine has not been resolved. Therefore, how these subunits interact cohesively *in situ*, is yet to be seen.

Here, we resolve the intact *in situ* architecture of a Tad pilus system–the *Cc*Tad pilus machine, by cryogenic electron tomography (cryo-ET) with subtomogram averaging. This approach is complemented by integrative modelling to elucidate the spatial arrangement of all core components of the *Cc*Tad pilus machine. Our data suggest that the OM pore complex has a novel composition, consisting of the secretin protein CpaC and the lipoprotein CpaD, but lacking CpaO, the TadD pilotin ortholog. Moreover, 3D classification reveals multiple distinct intermediates of the Tad pilus machine, allowing us to propose a stepwise assembly mechanism. Taken together, these results point to a refined architectural model for the Tad pilus system in *C. crescentus*.

## Results

### Identification and characterization of C. crescentus strains to maximize Tad pilus machine production for in situ structural studies

To achieve high-resolution *in situ* structures of the *Cc*Tad machine by cryo-ET and subtomogram averaging, it is necessary to image abundant copies of these machines within cellular tomograms. While the polar localization of the Tad pilus machine in *C. crescentus* presents an advantage for identifying and imaging the *Cc*Tad machine, there are nonetheless other challenges to overcome. First, *C. crescentus* cells exhibit a dimorphic life cycle wherein each cell division produces two phenotypically and morphologically distinct cell types: a non-motile, reproductive stalked cell and a motile, non-reproductive swarmer cell^20^. Tad pili are produced only by swarmer cells and are located exclusively at the flagellated pole of these cells. Thus, in a mixed population of *C. crescentus*, only a subset of cells are pilus-producing swarmer cells. Methods exist to synchronize *C. crescentus* cell populations to enrich for swarmer cells, but even in synchronized populations of the standard laboratory strain NA1000, only ∼20-40% of swarmer cells exhibit pili at any given time^27–29^. Moreover, these cells produce an average of only two pili per cell (Fig. 1A)^27^. Based on these factors, it is not feasible to resolve a high-resolution structure of the *C. crescentus* Tad pilus machine by cryo-ET and subtomogram averaging using the standard laboratory strain NA1000.To overcome these limitations, we utilized the *C. crescentus* bNY30a background, a hyperpiliated derivative of the *C. crescentus* CB13b1a strain, which produces many more pili per cell than NA1000^18,30^. This strain, which we hereafter refer to as *Cc*Hyp (for hyperpiliated), produces pili that extend and retract over time, indicating that they are functional (Fig. 1B, Movie S1).

**Figure 1:**
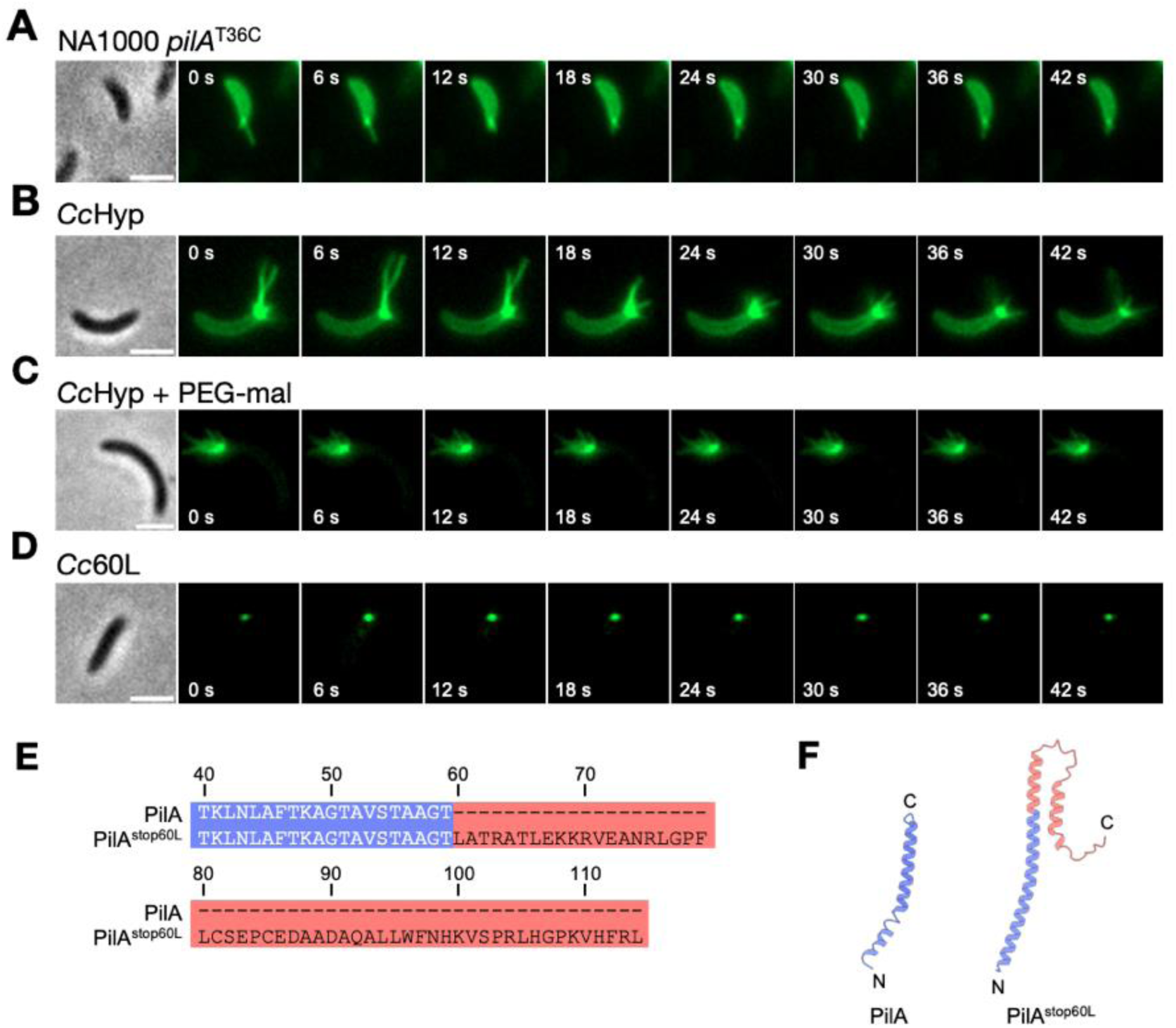
Identification of *C. crescentus* strains suitable for *in situ* structural determination of the Tad pilus machine. (**A-D**) Representative time-lapse images of the indicated strains labelled with AF488-maleimide (green), which reacts with the engineered cysteine residue in PilA^T36C^ enabling visualization of pili. In panel C, the strain was co-incubated with both AF488-maleimide and PEG5000-mal to label pili as well as artificially block pilus retraction. A representative phase contrast image of each time series is shown on the left. Scale bars, 2 μm. *Cc*Hyp: bNY30a Δ*hfsDAB pilA1*^T36C^; *Cc*60L: bNY30a Δ*hfsDAB pilA1*^T36C, stop60L^. (**E**) Alignment of the native PilA sequence with the stop60L variant of PilA (PilA^stop60L^). In PilA^stop60L^, the stop codon has been mutated to leucine, leading to the addition of 55 amino acids to the C terminus of the native PilA sequence. The amino acid numbers are indicated above the sequence. (**F**) Comparison of the structure of mature PilA (PDB 8U1K) to PilA^stop60L^ (predicted using AlphaFold3), depicting the effect of the 55 amino acid addition to the C terminus of PilA^stop60L^. Colouring corresponds to panel E.

To restrict the potential effect of pilus dynamics on our ability to image Tad machines *in situ*, we pursued two strategies. First, we artificially blocked pilus retraction in *Cc*Hyp by taking advantage of the engineered cysteine residue in PilA, which spontaneously reacts with the polymer methoxy-polyethylene glycol-maleimide with an average molecular weight of 5000 Da (PEG5000-mal)^31^. Upon reaction, the addition of bulky PEG5000-mal adducts to the pilus fibers prevents their retraction back into the cell, thus rendering these pili ‘frozen in time’ (Fig. 1C). This effect is likely due to steric exclusion of the pilus from passage through the OM secretin channel during retraction^27^. Our second strategy involved the use of a previously unpublished retraction-deficient variant of *Cc*Hyp, originally isolated during a screen for mutants that exhibited resistance to the pilus-dependent phage ΦCb5^18^. The mutation that produces this phenotype is in *pilA*, where the stop codon has been mutated to a leucine (*pilA^stop60L^*). This retraction-deficient strain, which we hereafter refer to as *Cc*60L, produces pili that appear as a single polar punctum rather than as extended fibers, likely due to an extremely short average length (Fig. 1D). Moreover, cells in the *Cc*60L population do not exhibit cell body fluorescence (Fig. 1D, S1) as observed for cells that are capable of normal pilus extension and retraction (Fig. 1A,B), indicating that pili produced by *Cc*60L are not retracted back into the cell after being fluorescently labeled outside^27^. At the protein level, the *pilA^stop60L^* mutation extends the length of PilA by 55 amino acids (Fig. 1E). This more than doubles the length of mature PilA in this mutant, increasing the length of its native α-helix, and adding a largely unstructured and presumably flexible region to its C terminus (Fig. 1F). Consequently, we hypothesize that the pilus phenotype of *Cc*60L results from early abortion of pilus extension combined with lack of pilus retraction, due to incompatible pilus filament geometry and/or steric blockage of the pilus filament within the OM secretin pore. Ultimately, both the above strategies yield cells that produce pili that cannot be retracted, either due to chemical modification of the pilus filament after extension (*Cc*Hyp + PEG5000-mal) or genetic mutation of PilA (*Cc*60L).

### Visualising the Tad pili of Cc60L and CcHyp in situ by cryo-ET

Having identified mutant strains and treatment strategies that improve Tad pilus abundance and visualization, we next examined their phenotypes by cryo-ET. Cryo-EM grids were prepared from *Cc*60L cells and from *Cc*Hyp cells treated with PEG5000-mal. These cells were grown to mid-exponential phase, when Tad pili are most abundant, and data were collected from cell poles using the recently developed parallel cryo-electron tomography (PACE-tomo) method^32^, enabling cryo-ET data acquisition in a high-throughput manner (> 200 tilt series per day)^23,33,34^.Reconstruction of 3D tomograms revealed striking phenotypic differences between the two strains (Fig. 2A-D). Among the 341 reconstructed tomograms of PEG5000-mal-treated *Cc*Hyp, only 128 cells contained visible pili (Fig. 2G). The piliated *Cc*Hyp cells typically displayed multiple filaments extending from the cell pole (Fig. 2A, B). The observed length of the retraction blocked *Cc*Hyp filaments often surpassed the 500 nm average (Fig. 2A, B, H; note: the pilus length in these cells frequently exceeded the tomographic field of view). Meanwhile, the corresponding *Cc*Tad machines typically spanned the entire periplasmic space, zipping the OM and IM closely together near the cell pole, with a periplasmic width of ∼ 33 nm at sites of piliated *Cc*Tad machines (Fig. 2I). In contrast, sites containing *Cc*Tad machines without pili had a larger average periplasmic width of ∼ 53 nm at the cell pole (Fig. 2I).

**Figure 2:**
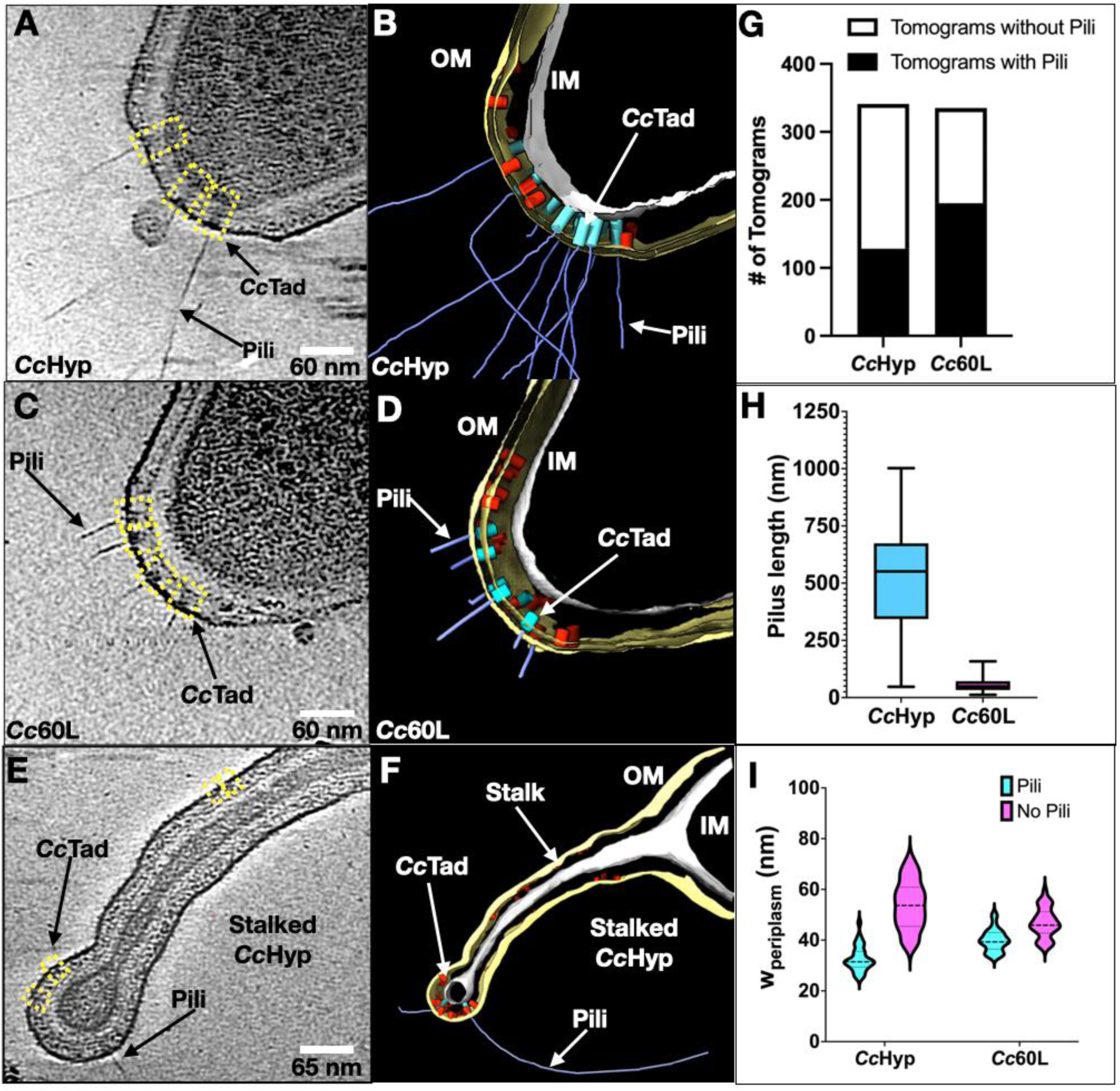
Cryo-ET reveals piliated *C. crescentus* Tad machines within the bacterial cell envelope. **(A)** A tomographic slice of a PEG5000-mal-treated *Cc*Hyp cell pole. Yellow boxes indicate the Tad pilus machines observed in the tomographic slice. **(B)** 3D segmentation of the tomogram shown in (A). The OM is colored yellow, the IM is colored grey, the piliated *Cc*Tad machines are colored cyan, the non-piliated *Cc*Tad machines are colored red, and Tad pili are colored blue. **(C)** A tomographic slice of a *Cc*60L cell pole. **(D)** 3D segmentation of the tomogram shown in (C). (**E)** A tomographic slice of a representative stalked PEG5000-mal-treated *Cc*Hyp cell, focused on the stalk region. (**F)** 3D segmentation of the tomogram shown in (E). (**G**) Quantification of tomograms containing pili in the *Cc*Hyp and *Cc*60L strains, respectively. (**H)** Measurement of the lengths of pili (N=30) from representative tomograms of *Cc*Hyp and *Cc*60L cells, respectively. (**I)** In cyan, violin plots showing periplasmic widths measured at the sites of piliated *Cc*Tad machines (N=30) at the cell poles of *Cc*Hyp and *Cc*60L cells, respectively. In magenta, the measured periplasmic widths of non-piliated *Cc*Hyp and *Cc*60L tomograms, respectively (N=30).

Conversely, a higher proportion of *Cc*60L cells were piliated (Fig. 2G), and piliated *Cc*60L cells typically produced slightly more Tad filaments per cell (752 pili in 195 piliated *Cc*60L cells; ∼3.86 pili/piliated cell) compared to PEG5000-mal-treated *Cc*Hyp cells (459 pili in 128 piliated cells; ∼3.59 pili/piliated cell). However, pili produced by *Cc*60L were much shorter in length, ranging from 10-150 nm (Fig. 2C, D, H). This phenotype is consistent with the appearance of pili as a single punctum in our fluorescently labeled images of the *Cc*60L strain (Fig. 1D, S1). Intriguingly, the Tad machines in *Cc*60L tomograms were typically smaller and predominantly anchored to the OM, without connection to the IM (Fig. 2D). Thus, the cell poles of piliated *Cc*60L cells typically had a wider periplasmic spacing of ∼ 40 nm, greater than in the piliated cell poles of *Cc*Hyp (Fig. 2I).

Stalked *C. crescentus* cells were seen in the populations of both strains. Stalked cells displayed fewer piliated machines, although incomplete Tad complexes were frequently observed at the tips of stalks, and along their lengths (Fig. 2E, F). Given the known absence of pili in stalked cells, we propose that these incomplete Tad machines are “relics” that persist within the stalked compartment after pilus activity has ceased at the stalked pole.

Altogether, our results show that while *Cc*60L cells provide abundant Tad pilus machines for *in situ* structure determination, especially of the OM complex, the PEG5000-mal-treated *Cc*Hyp cells yield a higher proportion of fully assembled, envelope-spanning pilus machines.

### Subtomogram averaging and 3D classification reveal the architectures of the Tad pilus machine at different stages of assembly

To further investigate the architectural details of the Tad pilus machine, 9,972 non-piliated and 1,211 piliated *Cc*Tad particles were manually picked from 675 tomograms of *Cc*60L and PEG5000-mal-treated *Cc*Hyp cells. To solve the structure of the non-piliated *Cc*Tad machine, we removed ∼ 40% junk particles by 3D classification. The initial subtomogram averaged structure revealed a prominent ring-like density anchored to the OM, resembling the secretin pore described in other type IV pilus systems^35,36^ (Fig. S2A). However, despite intact, envelope-spanning machines being visible in individual tomograms (Fig. 2A, B), no density could be seen in the lower periplasm, or at the IM. This could suggest a high degree of particle heterogeneity near the IM region, consistent with the observed variation in periplasmic widths (Fig. 2I).

To dissect this structural heterogeneity, we performed extensive 3D classification on the non-piliated *Cc*Tad machine particles. Multiple secretin-containing class averages were identified, consistent with distinct assembly intermediates of the Tad pilus machine (Fig. 3A-D). The first class contained only an OM-associated ring-shaped density (Fig. 3A), similar to the initial global average (Fig. S2A). A second class displayed additional density linking the base of the secretin to the peptidoglycan (PG) layer, suggesting recruitment of a periplasmic component at an early assembly stage (Fig. 3B; red arrow). A third class extended further below the now PG-bound secretin (Fig. 3C), forming a conduit in the periplasm that likely represents the alignment subcomplex. The IM was more clearly resolved in the fourth class, with additional densities in the cytoplasm, consistent with the fully assembled Tad pilus machine (Fig. 3D; red arrow). Notably, these fully assembled complexes represented fewer than 10% of all Tad pilus machine particles, whereas the majority (>80%) corresponded to the OM-associated intermediates (Fig. 3 A, B).

**Figure 3:**
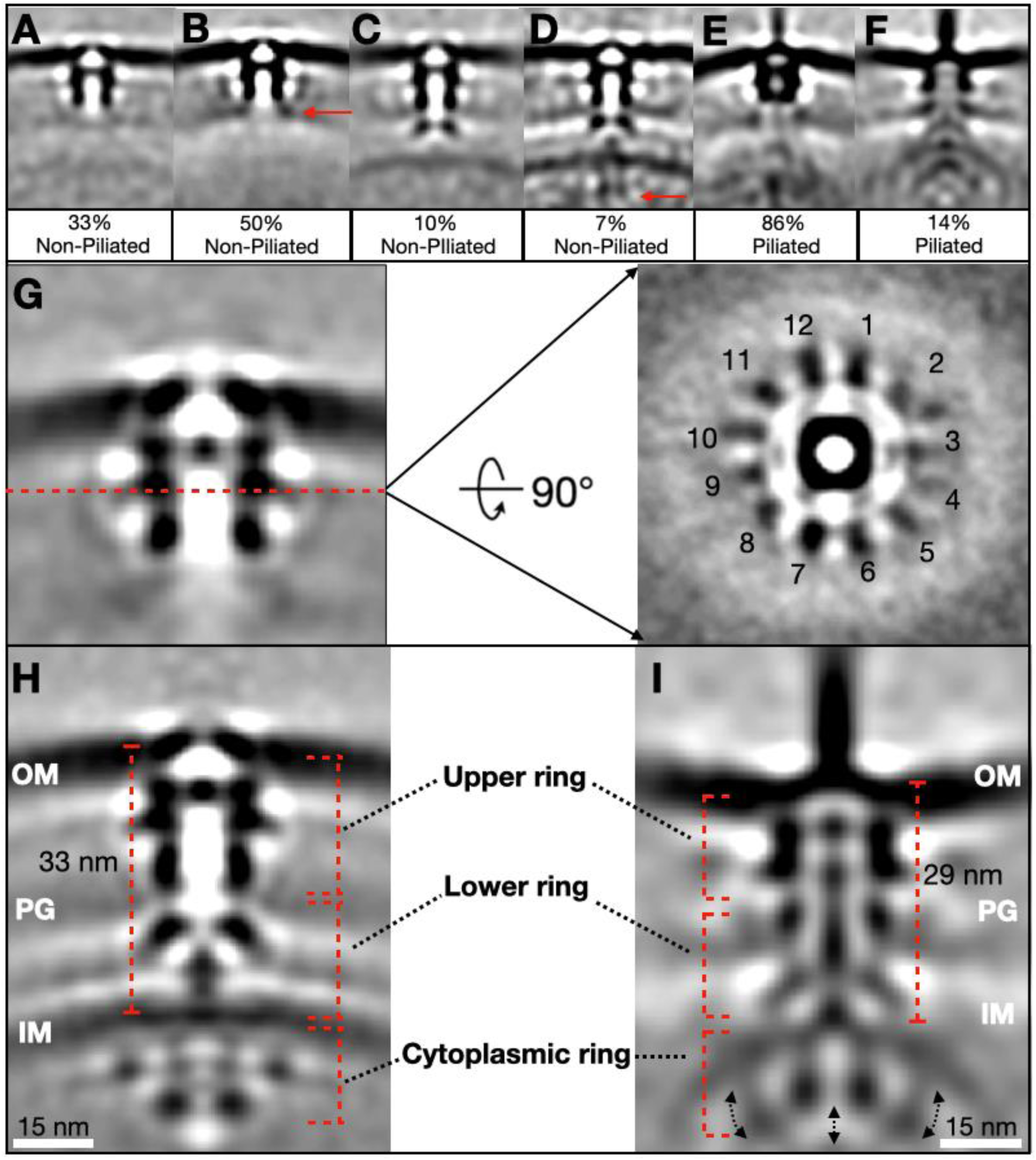
Subtomogram averaging and 3D classification reveal the *Cc*Tad machinery in different assembly intermediate states. **(A-F)** Central slices of class averages of non-piliated (A-D) and piliated (E-F) Tad machines. **(G)** On the left, a central slice of a focus-refined subtomogram-averaged density for the outer-membrane region of the non-piliated Tad machine. A cross-sectional view of the plane indicated by the red dashed line is shown on the right. **(H)** A central slice of a subtomogram-averaged structure (c6-symmetry expansion) of the non-piliated Tad machine. **(I)** A central slice of a subtomogram-averaged structure (c6-symmetry expansion) of the piliated Tad machine.

Meanwhile, the piliated *Cc*Tad machine particles were sorted into two primary classes through 3D classification (Fig. 3E, F). In one class, the pilus filament protruded through the OM-associated secretin, with no pilus density visible below the OM (Fig. 3E). The large majority of the particles in this class came from the *Cc*60L dataset, and were similar to the machines visible in the tomograms of this strain (Fig. 2C, D; Fig. S2B). The second class (Fig. 3F) resembled the fully assembled *Cc*Tad machine as seen in the non-piliated state (Fig 3D), but with a pilus filament protruding from the OM (Fig. 3F). This class likely represents intact Tad machines captured in the process of elaborating a pilus. As expected based on the individual tomograms (Fig. 2A, B), the majority (> 90%) of these piliated machines came from the PEG-mal blocked *Cc*Hyp cells. Together, these 3-D classification results reveal that the *Cc*Tad pilus machine adopts structurally heterogeneous states *in situ*. The diversity of class averages is consistent with a machine that undergoes dynamic assembly and disassembly. The abundance of partially assembled complexes likely reflects the transient nature of the Tad pilus in *C. crescentus*, due to cell-cycle coupled transcriptional control of *cpa* genes, which ensures that pili are only produced by swarmer cells at a specific stage to aid in surface sensing and initial attachment.

### Focused refinement reveals architectural differences between the non-piliated and piliated states

The large number of OM secretin-containing particles enabled focused refinement in the vicinity of the OM region. Rotational alignment revealed a c6-symmetric inner tube surrounded by spoke-like densities with 12-fold symmetry that extended to contact the PG layer (Fig. 3G). The observed arrangement of 6- and 12-fold symmetries are in line with the recently reported *P. aeruginosa* Tad secretin structure with 13- or 14-fold symmetry, as determined by single-particle cryo-EM^37^. We thus applied c6 symmetry expansion to our dataset, effectively increasing the particle number by a factor of six. This strategy enhanced map quality, enabling us to resolve structural features in the fully assembled machines in both the piliated and non-piliated states, to 4-5 nm resolution (Fig. 3H, I; Fig. S3).

Comparison of the two states clarified structural differences in both the machine and the surrounding cell envelope (Fig. 3H, I). In both states, the machine could be separated into three sections: the upper and lower periplasmic rings (spanning the OM, periplasm, and IM), and a cytoplasmic ring. The periplasmic spacing was larger in the non-piliated machine (∼ 33 nm) than in the piliated structure (∼ 29 nm), suggesting local remodelling of the cell envelope based on the state of the *Cc*Tad complex. In the non-piliated state, both the OM and IM were slightly convex in the vicinity of the Tad complex (Fig. 3H). Meanwhile, in the cytoplasmic ring, a central bell-shaped density was observed in association with the cytoplasmic leaflet of the IM, surrounded by smaller knob-shaped densities. In contrast, in the piliated state, the OM showed a concave curvature, whereas the IM exhibited pronounced convex bending (Fig. 3I). This bending was correlated with the downward movement of the knob-like densities in the cytoplasmic ring (Fig 3I, black arrows), now in contact with the central bell-shaped structure from below. Together, these conformational changes appear to drive the pronounced local curvature of the IM observed in the piliated state.

### Integrative modelling of the Tad pilus machine

To investigate the molecular basis of *Cc*Tad pilus assembly, we next fitted solved structures and AlphaFold3-predicted models of known Tad components into the focus-refined densities. The cylindrical, tunnel-shaped density in the OM region resembled secretins from other T4P systems^35–38^ (Fig. 4A, B) and was therefore assigned to CpaC, the *Cc*Tad secretin^23,34,39^. CpaC was predicted to adopt a classical secretin-like fold with a central β-barrel channel and upper-lip composed of an anti-parallel β-hairpin motif. Two globular domains localize below the secretin tunnel, including a classical secretin-N3-domain followed by an N-terminal β-sandwich VirB9-like domain^40^. The c12-symmetric dodecameric complex of CpaC fits well into the tunnel-shaped densities (average map value, 0.75). The upper lip remodels the OM at the secretin entrance in the extracellular space.

**Figure 4:**
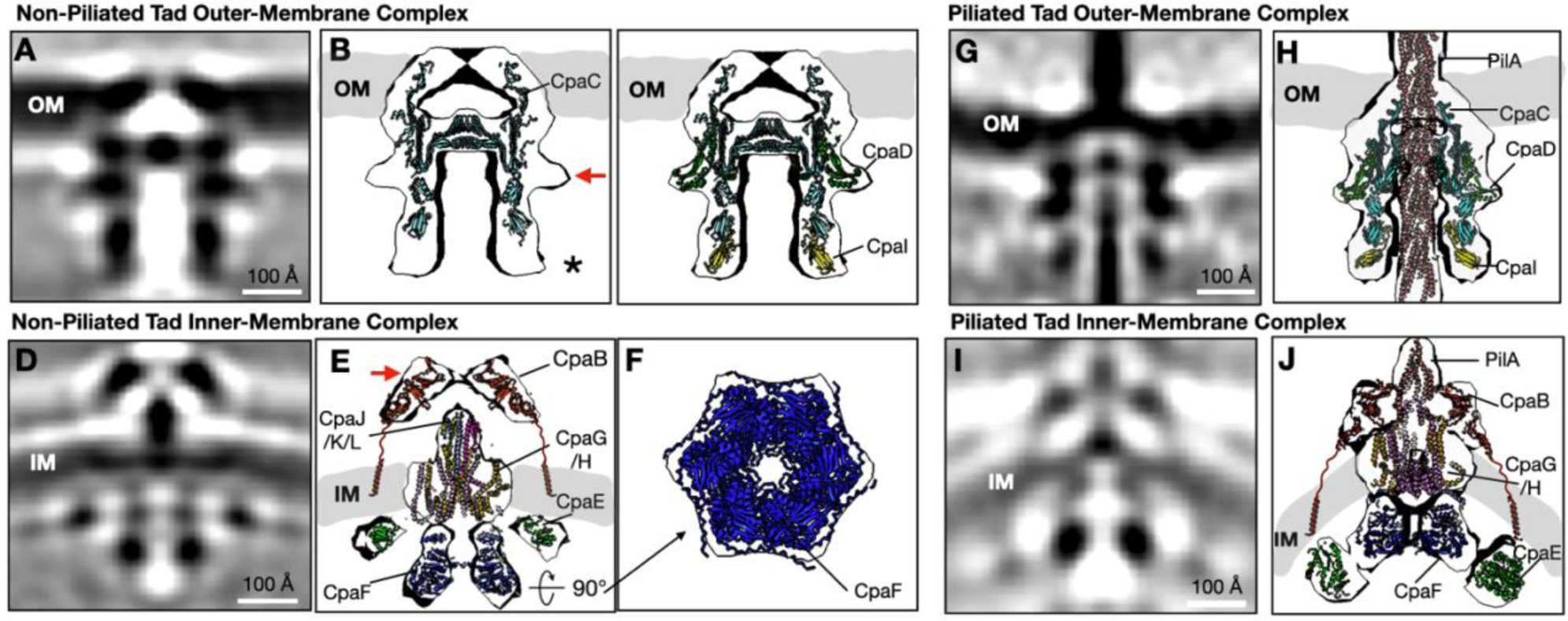
Focus refinement and integrative modelling reveals the architecture of the *Cc*Tad machine in piliated and non-piliated states. **(A)** A central slice of the focus-refined subtomogram-averaged map near the OM region of the non-piliated Tad pilus machine. (**B)** AlphaFold3-predicted models of CpaC fitted into the subtomogram-averaged map shown in (A). **(C)** AlphaFold3 predicted models of CpaC, CpaD and CpaI fitted into the subtomogram-averaged map shown in (A). **(D)** A central slice of the focus-refined subtomogram-averaged map near the inner-membrane region of the non-piliated Tad pilus machine. **(E)** AlphaFold3-predicted models of CpaB, CpaG, CpaH, CpaJ, CpaK, the pilin-like domain of CpaL, and CpaE, along with the Cryo-EM structure of CpaF, were fitted into the subtomogram-averaged map shown in (D). The flexible linkers between the transmembrane helices and globular domains of CpaB were drawn as squiggle lines. **(F)** Bottom view of the structure shown in (E). **(G)** A central slice of the focus-refined subtomogram-averaged map near the outer-membrane region of the piliated Tad pilus machine. **(H)** AlphaFold3-predicted models of CpaC, CpaD, CpaI and the Cryo-EM structure of the pilus were fitted into the subtomogram-averaged map shown in (G). **(I)** A central slice of the focus-refined subtomogram-averaged map near the inner-membrane region of the piliated Tad pilus machine. **(J)** AlphaFold3-predicted models of CpaB, CpaG, CpaH, CpaF and CpaE fit into the map shown in (I). The flexible linkers between the transmembrane helices and globular domains of CpaB were drawn as squiggle lines.

Two additional densities in direct contact with CpaC were consistently observed: a spoke-like belt encircling the β-barrel (Fig. 4B, red arrow) and another small PG-associated density located just beneath CpaC (Fig. 4B, black asterisk). CCNA_03033 (CpaO), previously proposed as the *Cc*Tad pilotin, was initially considered for the spoke-shaped belt density^41^. However, the AlphaFold3 model of CpaO in complex with CpaC indicated that its horseshoe-shaped, α-helix-rich structure failed to fit into the subtomogram-averaged density (Fig. S4). Consistent with this, deletion of *cpaO* did not impair pilus production or pilus-dependent phage sensitivity, indicating that CpaO is not required for pilus biosynthesis in *C. crescentus* (Fig. S5). However, the 225-amino-acid OM lipoprotein CpaD, encoded immediately downstream of *cpaC*, is required for Tad biogenesis in *C. crescentus*^23^, and is predicted to adopt an N-terminal OmpA-like periplasmic fold capable of binding PG, suggesting it may anchor the secretin to the PG layer^42^. Consistent with this hypothesis, AlphaFold3 multimer predictions identified an interaction interface between the CpaD and CpaC monomers (Fig. S6A), and an AlphaFold3-predicted CpaD dodecamer complex was in excellent agreement with the spoke-like density that wraps around the secretin (Fig. 4C).

The second unexplained PG-associated density, located just below CpaC (Fig. 4B, black asterisk), was assigned to the Tad component CpaI, a periplasmic protein of unknown function, located within the *cpa* locus and required for pilus biosynthesis^37,43,44^. CpaI is predicted to adopt a fold similar to the N-terminal domain of the *P. aeruginosa* Tad secretin RcpA, indicative of a functional association with the secretin^37^. The AlphaFold3-predicted structure of CpaI revealed a globular β-sandwich fold with a C-terminal solvent-exposed β-sheet comprised of three anti-parallel β-strands. AlphaFold3 multimer predictions further suggested that the CpaI C-terminal β-sheet packs against the N-terminal β-sandwich of the CpaC secretin in a manner reminiscent of donor-strand complementation (Fig. S7A, B, D), and the CpaI-CpaC complex fit well into the subtomogram averaged density (average map value, 0.9; Fig. 4C).

A remaining challenge was to understand how the OM-associated complexes connect with those anchored to the IM. Focused refinement of the regions near the IM in the non-piliated state further revealed several prominent densities (Fig. 4D, E). A large cashew-shaped periplasmic density above the IM likely represents the alignment subcomplex (Fig. 4E, red arrow). It was thus assigned to CpaB, which is predicted to contain two globular C-terminal domains that form an oblong structure, which fit well into the cashew-shaped density (average map value, 0.3). In addition, CpaB has an N-terminal transmembrane helix that anchors it to the IM^23^. Notably, the C-terminal residues of CpaB form an extended C-terminal β hairpin motif that is predicted to pack against the N-terminal β-sandwich domain of CpaI (Fig. S7C, D). These data suggest that CpaI acts as a bridge between the OM- and IM-associated Tad subcomplexes.

The proteins CpaG and CpaH are orthologous to the platform proteins found in other T4P systems^16^. CpaG and CpaH were predicted to form heterodimers (Fig. S6C)^26,37^, and a trimer of their heterodimeric complex fit well into the base of the central cone-shaped density within the IM. However, a small unassigned density remained at the tip of the cone. We hypothesized that it could correspond to the minor pilin-like proteins CpaJ, CpaK and CpaL, which are required for pilus production in *C. crescentus*^44^. CpaJ and CpaK are predicted to adopt similar structures to the T4aP and T4bP pilins^11^, and are thought to incorporate into the pilus tip in complex with the minor pilin-like protein CpaL, prior to pilus extension^29^. CpaL contains two distinct domains: a pilin-like domain^29^ and a C-terminal von Willebrand factor A (vWA) domain. We propose that CpaJ, CpaK and CpaL sit at the base of the Tad machine within the periplasmic space prior to pilus assembly, comparable to the role of the T4aP minor pilins in *P. aeruginosa*^36^. Indeed, the AlphaFold3-predicted complex of CpaJ, CpaK, the pilin-like domain of CpaL, and a trimer of CpaG-CpaH heterodimers (Fig. S6E) fit well into the complete density (Fig. 4E), reinforcing this hypothesis. However, no density was observed for the vWA domain of CpaL in our subtomogram averaged map, likely due to its flexibility relative to the pilin domain.

On the cytoplasmic side of the IM, the cryo-EM structure of the hexameric motor ATPase CpaF was docked into the central bell-shaped cytoplasmic density (Fig. 4E, F). The ring-shaped densities surrounding CpaF were fitted with six dimers of the ParA/MinD ATPase family protein CpaE, which is a ubiquitous component of the Tad pilus system, responsible for the proper localization of the secretin CpaC^11,39^. Structural studies of the CpaE homolog in *Eubacterium rectale*, TadZ, have revealed that it contains an atypical ATPase domain with a deviant Walker-A motif capable of binding ATP for dimerization, alongside an atypical receiver (REC) domain^45^. *C. crescentus* CpaE has a similar architecture and is also predicted to assemble as a homodimer (Fig. S6D)^11^. Additionally, CpaE has a C-terminal amphipathic helix that allows it to localize to the IM, like other proteins from the same family^39^. Taken together, the arrangement of these cytoplasmic *Cc*Tad components in the subtomogram averaged map is consistent with the known biology of these proteins, as well as with AlphaFold3-predicted interactions between CpaE and CpaF (Fig. S6B).

Subtomogram averaging of piliated particles produced maps of comparable quality to those of the non-piliated machines (Fig. 4G-J). In the OM region, the pilus filament was well-resolved as it traversed through the secretin pore complex, whose densities were assigned to CpaC, CpaD, and CpaI (Fig. 4G, H). The IM complex retained the same overall organization as was observed in the non-piliated state, with CpaB, CpaG/H, CpaF, and CpaE fitting into their corresponding densities (Fig. 4I-J). The two main distinctions were the presence of the pilus filament passing through the central channel of the machine, and the large conformational changes observed near the IM region (Fig. 4I, J). Specifically, in the piliated state, the cytoplasmic ATPase motor CpaF moves closer to the IM to interact with the platform proteins CpaG/H, while the IM-associated CpaE moves downward to be in contact with the base of the CpaF hexamer (Fig. 4J). The interaction between CpaE dimers is likely mediated by adjacent tandem CpaE REC domains. Meanwhile, although CpaB appears to be in contact with CpaE in our model (Fig. 4E, J; Fig. 5A, B, E, F), there is no direct evidence that supports their interaction.

**Figure 5:**
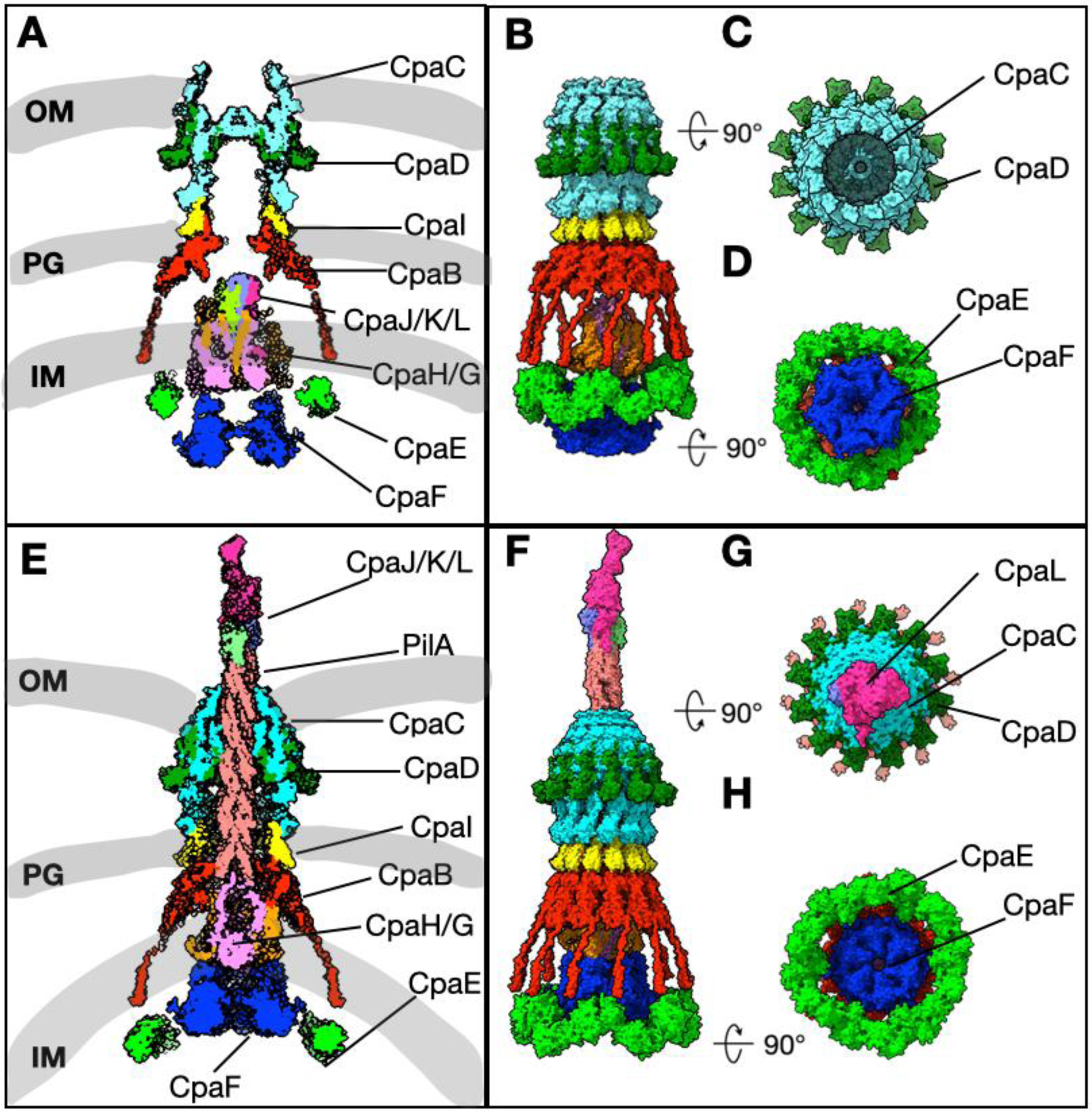
Architectural model of the *Cc*Tad pilus machine in the non-piliated and piliated states. **(A), (B)** An architectural model of the non-piliated *Cc*Tad pilus machine. **(C), (D)** Top and bottom views of the non-piliated *Cc*Tad pilus machine. **(E), (F)** An architectural model of the piliated *Cc*Tad pilus machine. **(F), (G)** Top and bottom views of the piliated *Cc*Tad pilus machine.

Taken together, *in situ* cryo-ET enabled fitting of cryo-EM solved structures and AlphaFold models of all *Cc*Tad components into the subtomogram averaged maps, resulting in proposed architectural models of the *Cc*Tad machine in both the non-piliated and piliated states (Figs. 4 and 5).

## Discussion

The Tad pilus is part of the TFF superfamily, a large and evolutionarily ancient group of envelope-spanning nanomachines that includes both bacterial and archaeal systems involved in motility, adhesion, and secretion^2,16^. Previous studies have inferred models of the Tad pilus machine largely from genetic analyses and homology-based comparison to the T4aP and T4bP^11,12,41^ systems.. While these analyses have provided a conceptual framework for Tad pilus assembly and function, they offer only a partial view of its biology, and often obscure its distinct evolutionary trajectory within the TFF superfamily. Furthermore, high-resolution structures of isolated Tad components, including *P. aeruginosa* RcpA and RcpC, which form a membrane-spanning pore complex, and *C. crescentus* CpaF, an ATPase driving pilus extension and retraction, provide valuable insights into the domain architecture and function of these key Tad components^12,16,17,26,37^. However, the *in situ* arrangement of these elements within an assembled Tad machine have not been visualized. As a result, the spatial organization and connectivity of Tad machine components has remained highly speculative. Our work presents the *in situ* architecture of a Tad pilus machine, resolved by cryo-ET and subtomogram averaging. Through an integrative modelling approach, we directly visualize how this nanomachine is organized across the *C. crescentus* cell envelope, providing a structural framework to interpret Tad systems within the broader context of the bacterial-archaeal TFF superfamily (Fig. 5).

Obtaining sufficient cryo-ET data to resolve the Tad pilus machine presents inherent challenges. In *C. crescentus*, the Tad pilus acts primarily as a dynamic surface-sensing appendage rather than as a stable adhesion structure^20,22,27^. It undergoes rapid cycles of extension and retraction that trigger holdfast synthesis upon surface contact, a critical step in the bacterium’s transition from a motile to a sessile lifestyle. The highly dynamic nature of the *C. crescentus* Tad pilus likely explains the scarcity of fully assembled machines in wild-type cells and the frequent observation of isolated secretin pores in our tomograms (Fig. 3A, B; Fig. S2). In contrast, Tad systems in other bacteria such as *A. actinomycetemcomitans* or *P. aeruginosa* are likely to be less dynamic as they are engaged in anchoring bacteria within the biofilm matrix. However, the dense, multicellular biofilms produced by these bacteria pose their own technical barriers for cryo-ET imaging, which require cryo–focused-ion-beam (cryo-FIB) milling to obtain vitrified, electron-transparent lamellae prior to cryo-ET data acquisition.

To overcome the low abundance and instability of the *C. crescentus* Tad machine, we combined genetic and chemical strategies to enrich for assembled complexes. The *Cc*60L mutant increased the number of observable Tad machines, while PEG5000-mal treatment of *Cc*Hyp cells further enhanced the proportion of extended pili by chemically blocking pilus retraction. Although both approaches could potentially introduce artifacts: PEG5000-mal through covalent linkage with the cysteine-modified PilA subunit, and the *pilA* mutation in *Cc*60L through structural alterations to the pilus filament, the Tad pilus machines embedded within the cell envelope are themselves unlikely to be affected by these extracellular perturbations. Together, these strategies provided a sufficiently large population of intact complexes for *in situ* structural analysis. Using a high-throughput data acquisition workflow with PACE-tomo^28,32^, we collected over 200 tilt series per day, substantially expanding the datasets and enabling higher-quality subtomogram averaging and stringent 3D classification. This comprehensive strategy allowed us to capture the structural diversity of the Tad machinery, revealing a continuum of distinct assembly intermediates that reflect the dynamic cycles of pilus extension, retraction, and turnover in *C. crescentus* in response to environmental stimuli^11,20,22,27^. Extensive 3D classification was critical for disentangling the structural heterogeneity of the datasets and for isolating distinct class averages corresponding to the non-piliated and piliated states. These class averages provided the foundation for focused refinement and integrative modelling, enabling a direct visualisation of the Tad machine’s molecular architecture *in situ.* The resulting model not only validates, but significantly advances previous conceptual frameworks of the Tad apparatus by resolving its subunit organisation within the cell envelope.

Earlier models proposed that the secretin CpaC associates with CpaO, based on homology to the Tad systems in *A. actinomycetemcomitans* and *P. aeruginosa,* where CpaO orthologs function as pilotins that assist secretin assembly and localisation^37,46^. In *C. crescentus*, CpaO is non-essential, and its predicted structure did not fit the corresponding densities in our cryo-ET maps (Fig. S4 and S5). Transcriptomic data further show that *cpaO* is expressed much later in the cell cycle than the genes encoding other OM components of the Tad system, supporting the conclusion that it does not participate in Tad machine assembly^47^. Instead, our modelling suggests that the lipoprotein CpaD forms a complex with CpaC, consistent with its predicted topology and localisation. Notably, the structure of the *P. aeruginosa* Tad secretin RcpA was solved without a CpaD ortholog, suggesting that this strategy is not universally conserved among the Tad systems^37^. Together, these findings demonstrate that in *C. crescentus*, the CpaC-CpaD complex constitutes the OM module of the Tad machine, anchoring the apparatus and providing a structural foundation for subsequent periplasmic and cytoplasmic assembly.

Beneath the OM-associated CpaC-CpaD complex, the small periplasmic protein CpaI is predicted to occupy a strategic position bridging the secretin and the IM–anchored alignment subcomplex (Fig. 4C; Fig. 5; Fig. 6). Through this interface, CpaI may stabilize the Tad machine across the PG layer and cell envelope during pilus extension and retraction. Comparative analyses indicate that Tad systems across species have evolved diverse mechanisms for connecting their OM and IM complexes, as many lack a clear CpaI ortholog^46^. For instance, in *P. aeruginosa* the secretin RcpA directly associates with the CpaB ortholog RcpC, a subunit of the alignment subcomplex^12^.

**Figure 6:**
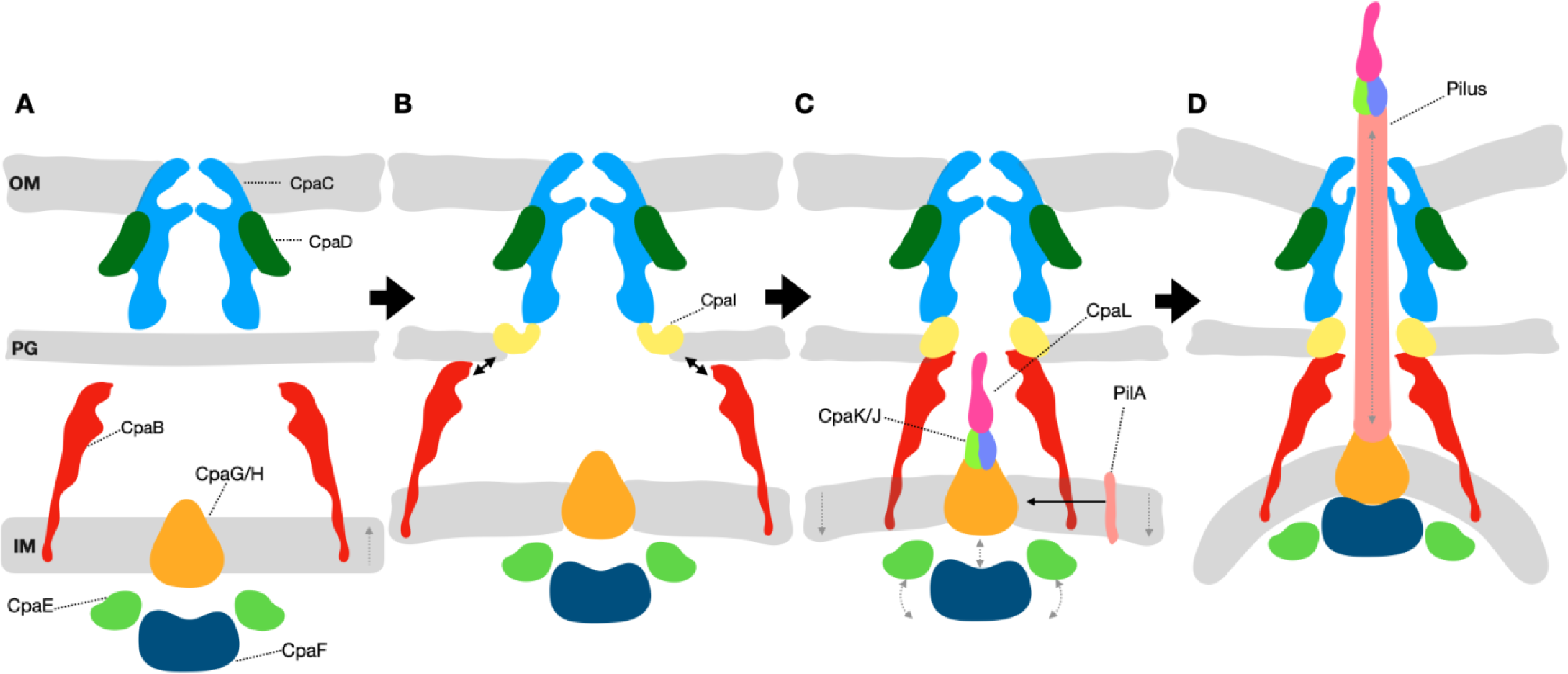
Proposed model of *Cc*Tad pilus assembly. **(A)** The secretin consisting of CpaC (light blue) and CpaD (green) is inserted into the OM. The assembly platform and motor subcomplex consisting of CpaG/H (orange), CpaE (light green) and CpaF (dark blue) are formed at the IM and the cytoplasm. The IM-associated CpaB monomers (red) in the periplasm are localized near the secretin. **(B)** In the periplasm, CpaI (yellow) is recruited to connect the OM-associated CpaC with PG as well as the IM-associated CpaB (red). **(C)** Stable assembly of the secretin and alignment subcomplex enables the recruitment of the motor subcomplex and the minor pilins CpaJ (lime green), CpaK (purple), and CpaL (cyan). (**D**) Full assembly of the nanomachine primes the polymerisation of the pilus filament using the major pilin PilA (pink) and drives the rapid cycles of pilus extension and retraction.

The alignment subcomplex, assembly platform and motor subunits in *C. crescentus* are formed by CpaB, CpaG/H, CpaE, and the ATPase CpaF, which assemble into a multi-ring structure that anchors the nanomachine to the IM and powers pilus dynamics. Transcriptional data for the *cpa* genes further suggests that the IM and OM subcomplexes assemble early, whereas *cpaI* is expressed later in the cell cycle^41^, consistent with a role for CpaI as a bridge that completes the connection between the two membrane-associated subcomplexes. These findings support a sequential assembly pathway for the *C. crescentus* Tad pilus machine (Fig. 6). As the secretin CpaC is integrated into the OM with CpaD (Fig 6A), IM-associated CpaB is localized near the secretin but is not yet linked with the OM-associated complex (Fig 6A). Next, CpaI is recruited near the PG layer (Fig. 6B) and acts as a bridge to connect the CpaB alignment subcomplex to the secretin, to form the framework of the Tad pilus machine spanning the entire bacterial cell envelope. Notably, CpaB appears to form a cage-like structure reminiscent of the PilN-PilO-PilP complex in the T4aP systems^36^, to help position the platform and motor subcomplex components (Fig 5A, B, E, F), prompting the further assembly of minor pilins CpaK and CpaJ as well as the Tad pilus tip CpaL (Fig. 6C)^29^. This might suggest a mechanism reminiscent to that of the minor pilins in the T4aP of *M. xanthus*, where they prime the assembly of the pilus^48^. Thereafter, PilA monomers embedded in the IM can be incorporated into the growing pilus filament, enabling its extension from the cell (Fig. 6C, D)^11^. This model thus defines how the Tad pilus machine assembles across the bacterial cell envelope, ultimately forming a continuous conduit linking the cytoplasmic motor to the extracellular pilus filament (Fig. 6).

The conformational differences observed between the non-piliated and piliated states, most notably in membrane curvature and the positioning of the motor ATPase CpaF, highlight the dynamic coupling between mechanical force generation and cell-envelope remodeling during pilus extension and retraction. Similar transitions in membrane curvature have been reported for other TFF systems, flagella, and type IV secretion systems^5,36,49–51^. While the functional consequences of membrane remodelling remain poorly understood in bacteria, in eukaryotic systems membrane curvature often plays key regulatory roles by recruiting proteins such as kinases to initiate signalling^52,53^. Thus, membrane remodelling events may not only accommodate mechanical strain imposed by force transmission but could also influence the assembly state and activity of cell-envelope-spanning nanomachines in bacteria.

In summary, we present the *in situ* architecture of the *C. crescentus* Tad pilus machine, resolved through a combination of bacterial genetics, high-throughput cryo-ET with subtomogram averaging, and molecular modelling. This work refines our understanding of the Tad machine organization in *C. crescentus* and provides a structural framework for comparing TFF systems across diverse bacterial and archaeal species. Future efforts in imaging Tad subunit-deletion mutants will validate their localisation within the nanomachine, and will aim to reveal additional assembly intermediates that may provide insight into motor dynamics. Further leveraging cryo-FIB milling will be key for revealing architectures of other Tad machines in their native states within biofilm matrices.

## Materials and Methods

### Bacterial strains, plasmids, and growth conditions

Bacterial strains, plasmids, and primers used in this study are listed in Table S1. *C. crescentus* strains were grown at 30 °C in peptone-yeast extract (PYE) medium^54^. Commercially available, chemically competent *Escherichia coli* DH5a (NEB5α, New England Biolabs) was used for plasmid construction and was grown at 37 °C in lysogeny broth (LB) supplemented with 25 μg/mL kanamycin (Kan), where appropriate, for plasmid maintenance.

Plasmids were transferred to *C. crescentus* by electroporation, as described previously^55^. Chromosomal mutations were made by double homologous recombination using pNPTS138-derived plasmids, as previously described^56^. Briefly, plasmids were introduced into *C. crescentus* by electroporation, then two-step recombination was performed using Kan resistance to select for single crossovers, followed by sucrose resistance to identify plasmid excision events. All mutants were validated by Sanger sequencing using primers targeting outside the region of recombination to confirm the presence of the mutation.

For construction of the pNPTS138-derived plasmids, ∼500-bp regions of DNA flanking either side of the desired mutation were amplified from *C. crescentus* NA1000 or CB13 genomic DNA, as appropriate. Upstream regions were amplified using upF and upR primers, and downstream regions were amplified using downF and downR primers (Table S1). upF and downR primers contained flanking sequences to facilitate insertion into pNPTS138 digested with EcoRV (New England Biolabs) by Gibson assembly (HiFi DNA Assembly Master Mix; New England Biolabs). Assembled plasmids were transformed into *E. coli* NEB5a (New England Biolabs), and clones with a positive insert were verified by Sanger sequencing (Table S1).

### Phage sensitivity assays

Phage sensitivity assays were performed using the pilus-specific phages ΦCbK and ΦCb5, as described previously^57^. Briefly, 400 μL of *C. crescentus* stationary phase culture was mixed with 4 mL of 0.75% (w/v) molten PYE top agar. The mixture was spread over a 1.5% (w/v) PYE agar plate and incubated at room temperature for 1 h to solidify. A tenfold serial dilution series of ΦCbK or ΦCb5 was prepared in PYE, and 2 μL of each dilution was spotted on the agar plate. The plates were grown for 2 days at 30 °C before imaging using a ChemiDoc MP (BioRad).

### Pilus labelling, blocking, and imaging

The pili of *C. crescentus* were labelled as described previously^31^. Briefly, 25 μg/mL of Alexa Fluor 488 C5 Maleimide (AF488-mal, ThermoFisher Scientific) was added to 100 μL of early exponential phase *C. crescentus* cell culture (OD_600_ = 0.1–0.3) and incubated for 5 min at room temperature. To artificially block pilus retraction, 500 μM of methoxy-polyethylene glycol maleimide with an average molecular weight of 5 kDa (PEG5000-mal, Sigma) was added to the *C. crescentus* cell culture immediately before the addition of 25 μg/mL AF488-mal. Labelled and/or blocked cells were collected by centrifugation at 5000 × g for 1 min, and were washed once with 100 μL of PYE to remove excess dye. The cell pellets were resuspended in 20 μL of PYE, one μL of which was spotted onto a 1% agarose PYE pad (SeaKem LE, Lonza Bioscience). The agarose pad was sandwiched between glass coverslips for imaging, which was performed using a Nikon TiE inverted fluorescence microscope with a Plan Apo 60× objective, a green fluorescent protein (GFP) filter cube, a Hamamatsu OrcaFlash 4.0 CCD camera, and Nikon NIS Elements imaging software. Images were analyzed using ImageJ software^58^.

### Preparation of cryo-EM grids

*C. crescentus Cc*Hyp and *Cc*60L cells were grown overnight on PYE agar plates at 30°C. Single colonies were later transferred into liquid PYE medium and serially diluted 5 times, then again cultured overnight at 30°C with shaking to reach early- to mid-exponential phase. For *Cc*60L cells, a dilution with an OD_600nm_ of 0.2 was mixed with 40% 10-nm BSA fiducial nanogold tracer (Aurion). For *Cc*Hyp cells, a dilution with an OD_600nm_ of 0.2 was mixed with 40% 10-nm BSA fiducial nanogold tracer and 500 μM of PEG5000-mal. The bacterial solutions were deposited on freshly glow-discharged 300 mesh holey carbon grids (C-Flat, Electron Microscopy Sciences) for 5 min. Grids were manually blotted with filter paper, then vitrified in liquid ethane using a custom gravity-driven plunger apparatus. The frozen grids were stored in liquid nitrogen until data acquisition.

### Cryo-ET data acquisition and tomogram reconstruction

The frozen grids were loaded into a 300-kV Titan Krios electron microscope (Thermo Fisher Scientific) equipped with a K3 Direct Electron Detector and BioQuantum energy filter (Gatan) at the McGill Facility for Electron Microscopy Research (FEMR). The PACEtomo^32^ script was used with the SerialEM software^28^ to collect multiple tilt series simultaneously with defocus values of approximately −4.8 μm, and a cumulative dose of ∼100 e−/Å covering angles from −48° to 48° (3° tilt step) or –60° to 60° (3° tilt step). Images were recorded at 42,000× magnification with an effective pixel size of 2.12 Å. All images were initially drift-corrected using MotionCor3^59^ and stacked with the IMOD software package^60^. Then, images were subsequently aligned in IMOD^60^ using 10-nm gold fiducial markers. Tomograms were reconstructed using TOMO3D^61^ into both the simultaneous iterative reconstruction technique (SIRT) and weighted back-projection (WBP) methods.

### Measurement of Tad pilus length and periplasmic spacing

Tomographic reconstructions were analysed to measure pilus length and periplasmic width. For randomly selected tomograms, coordinates corresponding to the boundaries and intermediate directional changes were recorded using IMOD. The Euclidean distance between the corresponding coordinates were calculated using the formula:

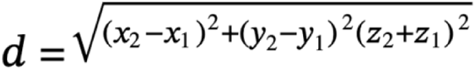

Statistical analysis and data visualization were performed using GraphPad Prism version 10.6.1.

### Subtomogram analysis

*Cc*Tad machines were manually picked in 6x binned SIRT tomograms for the best contrast-to-resolution ratio. The i3 software package was used for aligning and centering the particles and creating a subtomogram average using the WBP tomograms^62^. Multivariate statistical analysis (MSA)^63^ and 3D classification were used to remove junk particles from the picked dataset and visualize different 3D states. The remaining particles were aligned and centred again by using the i3 software package. Properly aligned machines were then brought to 4x binning and aligned further for higher resolution structures. Piliated and non-piliated particles were selected separately and their structures were therefore solved independently through extensive 3D classification. The focus-refined structure of the secretin pore complex was solved at 4x binning using the i3 package. Rotational alignment and 3D classification allowed for elucidation of the most probable structural symmetry, c6. Symmetry expansion was performed to expand the dataset and improve the subtomogram-averaged structure. These particles were used for the final global averages.

### Molecular modelling and visualization

Protein FASTA sequences for our model building were found on the NCBI protein database. The solved cryo-EM structure of CpaF (PDB ID: 8RKD) was used for fitting into our maps. We used the default settings of AlphaFold3^64^ to predict monomer and multimer structures of all other Tad pilus protein subunits to fit into the Cryo-ET maps generated in our study. Our Cryo-ET maps were exported to UCSF ChimeraX^65^ after inverting densities and adjusting to the correct pixel sizes. Model fitting was performed in UCSF ChimeraX using the “fit in map” function, which maximizes the average map value at atomic positions. For each atom within the map boundaries, the local density value was obtained by trilinear interpolation from the eight surrounding grid points of the map. The mean of these interpolated values represents the average map value, which was used as the fitting criterion. Segmentation of the tomogram pili and Tad machines was done using IMOD^60^. Cell membranes were segmented using MemBrain V2^66^.

## Data Availability

The sub-tomogram averaged structures of the *Cc*Tad secretin, the full non-piliated and piliated machines generated in this study have been deposited into the Electron Microscopy Data Band (EMDB) with the accession codes of EMD-73646, EMD-73615 and EMD-73632, respectively.

## Acknowledgements

We thank Dr. Kaustuv Basu at the McGill Facility for Electron Microscopy Research (FEMR) for their assistance with cryo-ET data collection. We thank Dr. Mike Strauss for access to a manual plunge freezing device and for PACE-tomo software installation and setup. We thank Mollee Ye for assistance in optimizing the cryo-ET workflows. We thank Velocity Hughes (Synthesis by Velocity, Malmö, Sweden) for editorial help with the manuscript. S.G is supported by an NSERC Discovery Grant (RGPIN-2024-04631), a Fonds de recherche du Québec – Santé (FRQS) Research Scholar (Junior 1) award (359456 & 376506), a McGill Centre de Recherche en Biologie Structurale (CRBS) Bluesky award, and a start-up fund by the McGill Faculty of Medicine and Health Sciences. J.I is supported by studentships awarded by the CRBS, Canadian Antimicrobial Resistance Network (CAN-AMR-NET) and the Antimicrobial Resistance Centre (AMR) McGill. G.B.W is supported by postdoctoral fellowships awarded by the Natural Sciences and Engineering Research Council of Canada (NSERC) and Fonds de recherche du Québec – Nature et Technologies (FRQNT); C.K.E was supported by a fellowship (1342962) from the National Science Foundation (NSF); Y.V.B is supported by the Canada 150 Research Chair in Bacterial Cell Biology and a grant from the National Institutes of Health (R35GM122556).

## Author contributions

SG, YVB, GBW and JI conceived the research; JI, RFW, GBW, CKE and SG performed the experiments; JI, GBW, RFW and SG wrote the manuscript. All authors contributed to the editing and revision of the manuscript.

